# Nanopore direct RNA sequencing detects differential expression between human cell populations

**DOI:** 10.1101/2020.08.02.232785

**Authors:** Josie Gleeson, Tracy A. Lane, Paul J Harrison, Wilfried Haerty, Michael B Clark

## Abstract

Accurately quantifying gene and isoform expression changes is essential to understanding cell functions, differentiation and disease. Therefore, a crucial requirement of RNA sequencing is identifying differential expression. The recent development of long-read direct RNA (dRNA) sequencing has the potential to overcome many limitations of short and long-read sequencing methods that require RNA fragmentation, cDNA synthesis or PCR. dRNA sequences native RNA and can encompass an entire RNA in a single read. However, its ability to identify differential gene and isoform expression in complex organisms is poorly characterised. Using a mixture of synthetic controls and human SH-SY5Y cell differentiation into neuron-like cells, we show that dRNA sequencing accurately quantifies RNA expression and identifies differential expression of genes and isoforms. We generated ∼4 million dRNA reads with a median length of 991 nt. On average, reads covered 74% of SH-SY5Y transcripts and 29% were full-length. Measurement of expression and fold changes between synthetic control RNAs confirmed accurate quantification of genes and isoforms. Differential expression of 231 genes, 291 isoforms, plus 27 isoform switches were detected between undifferentiated and differentiated SH-SY5Y cells and samples clustered by differentiation state at the gene and isoform level. Genes upregulated in neuron-like cells were associated with neurogenesis. We further identified >30,000 expressed transcripts including thousands of novel splice isoforms and transcriptional units. Our results establish the ability of dRNA sequencing to identify biologically relevant differences in gene and isoform expression and perform the key capabilities of expression profiling methodologies.

## Introduction

Cellular fates and functions are underpinned by the expression of protein-coding and non-coding genes into RNA (termed the transcriptome). The expression profiles of individual genes can vary in complex ways to regulate their functional outputs. Expression of genes can be switched on or off, increased or decreased, while the RNA products (isoforms) made from individual genes can also vary extensively. In humans >90% of protein-coding genes express multiple RNA isoforms via processes such as alternative transcriptional start sites, termination sites and splicing, greatly increasing the diversity of the transcriptome and proteome within cells (Pan et al. 2008; Wang et al. 2008). Expression of different genes and isoforms drive cellular differentiation programs, control cell and tissue functions and allow cells to respond to their environment (Melé et al. 2015; Roundtree and He 2016). However, aberrant expression contributes to various diseases including neurological disorders, autoimmune disorders and cancer (Emilsson et al. 2008; Lee and Young 2013; Sui et al. 2014).

Understanding the dynamic transcriptome requires techniques that can identify differential expression (DE) at both the gene (DGE) and transcript isoform (DTE) levels. Differential expression analysis enables comparisons between different tissues or conditions to identify genes that play a major role in determination of the phenotype. Short-read RNA sequencing (RNA-seq) methodologies are well validated for identifying differential gene expression but have limitations in identifying and quantifying both known and novel alternative isoforms (Steijger et al. 2013; Cali et al. 2019). This is exacerbated in complex mammalian transcriptomes which contain large numbers of highly similar transcript isoforms. A further limitation of short-read RNA-seq is the necessity to reverse transcribe RNA into cDNA and PCR amplify samples before sequencing, which introduces various biases (Aird et al. 2011).

Long-read sequencing techniques from Oxford Nanopore Technologies (ONT) and Pacific Biosciences have the potential to overcome many of these limitations (Sharon et al. 2013; Weirather et al. 2017). Long-read methods can sequence entire transcripts in a single read, potentially allowing the unambiguous identification of the expressed gene and isoform. However, initial long-read methods still required PCR and/or reverse transcription. Recently, ONT developed direct RNA sequencing (dRNA), the first long-read technique to sequence native RNA molecules (Garalde et al. 2018). dRNA doesn’t utilise any amplification or fragmentation steps and has the potential to quantify both genes and isoforms in an unbiased manner, while also characterising the RNA modifications and polyA tail on each RNA. Studies to date have used dRNA sequencing to catalogue known and novel transcripts in yeast (Jenjaroenpun et al. 2018), *C*.*elegans* (Roach et al. 2020; Li et al. 2020), *Arabidopsis* (Zhang et al. 2020) and human cell lines (Workman et al. 2019; Soneson et al. 2019); characterise polyA tail lengths of individual transcripts (Roach et al. 2020; Workman et al. 2019); identify allele specific gene and isoform expression (Workman et al. 2019); identify RNA base modifications (Workman et al. 2019; Garalde et al. 2018; Liu et al. 2019; Lorenz et al. 2019) and infer RNA structure (Stephenson et al. 2020). Maximum read lengths for dRNA were also much longer than for PCR-based long-read cDNA sequencing (Workman et al. 2019), demonstrating its potential to sequence long and complex RNA splice isoforms.

Identification of expression differences between samples is a standard requirement in the transcriptomics field. Therefore, dRNA sequencing needs to effectively quantify RNA and identify differentially expressed genes and isoforms to become a mainstream transcriptomics technique. The unbiased nature of dRNA sequencing should allow accurate quantification and has previously shown good performance on synthetic control RNAs of known abundance (Garalde et al. 2018; Sessegolo et al. 2019). However, the ability of dRNA sequencing to identify differential gene and isoform expression has largely been performed on model organisms with much simpler transcriptomes than found in mammals and/or based on expression fold changes without statistical analysis (Jenjaroenpun et al. 2018; Li et al. 2020; Zhang et al. 2020). Therefore, it remains essential to establish the effectiveness of dRNA sequencing to identify differential expression in complex organisms.

Here we use a combination of synthetic spike-in control RNAs and a neuroblastoma differentiation paradigm to demonstrate that dRNA sequencing accurately quantifies genes and isoforms and is able to identify biologically relevant differential gene and transcript isoform expression. We find that variance between samples is dominated by biological differences, allowing the identification of hundreds of DE genes and isoforms despite the lower throughput of dRNA sequencing. We further show that dRNA sequencing can identify differential transcript usage (DTU), where switching occurs between isoforms, often in the absence of overall changes to gene expression. Lastly, we demonstrate that dRNA sequencing can identify thousands of known and novel transcripts in SH-SY5Y (5Y) cells confirming the potential of dRNA sequencing to help fully decipher the complex transcriptome.

## Results

To examine the ability of dRNA sequencing to identify differential expression from a complex mammalian transcriptome, we utilised the well characterised differentiation of the human neuroblastoma SH-SY5Y (5Y) cell line into neuron-like cells. Native RNA from duplicate samples of undifferentiated and triplicate samples of differentiated 5Y cells were sequenced on an Oxford Nanopore MinION (**Figure 1A**). In addition, the RNA extracted from one differentiated sample was sequenced twice, producing a technical replicate to examine the variability due to library preparation and sequencing. Synthetic RNA ‘sequin’ spike-in controls (Hardwick et al. 2016) (see methods) were included in all samples to provide positive controls for gene and isoform identification, quantification and differential expression. Sequin RNAs vary in abundance over >4 orders of magnitude and come in two mixes, each mix contains the same synthetic RNA isoforms but their concentrations are offset by known amounts. Mix A was added to undifferentiated 5Y RNA, while Mix B was added to RNA from differentiated samples. The yeast calibration RNA was also included to enable initial quality control of sequencing.

**Figure 1.**
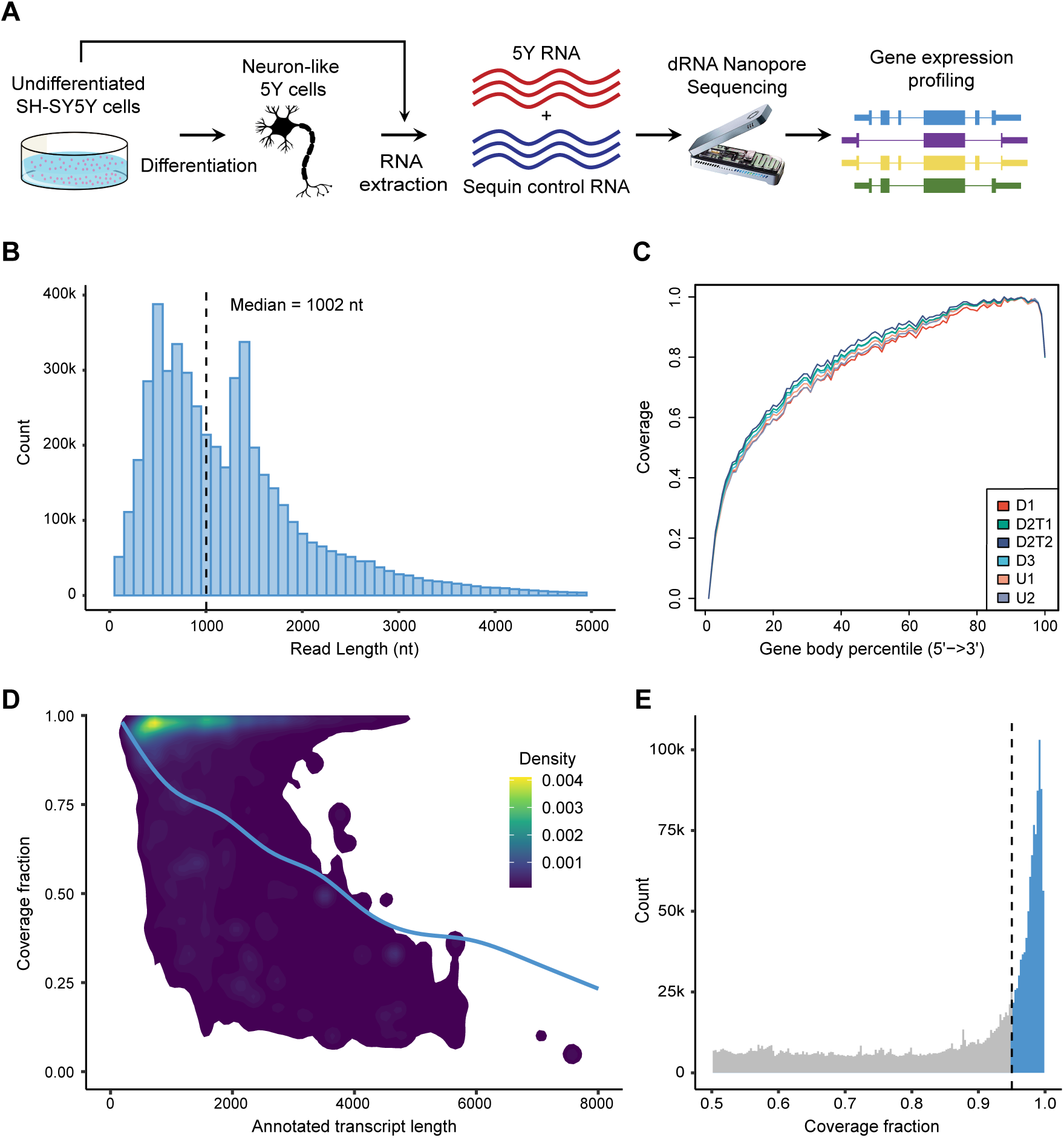
Length and coverage of direct RNA sequencing reads. **(A)** Experimental overview. Cultured SH-SY5Y cells were differentiated in triplicate and RNA extracted from both undifferentiated and differentiated cells. Native polyA purified SH-SY5Y RNA was combined with ‘sequin’ spike-in RNA, prepared for dRNA sequencing and sequenced on an Oxford Nanopore MinION. Sequenced reads were base-called, filtered and analysed to identify and quantify genes and transcript isoforms and their differential expression between undifferentiated and differentiated cells. **(B)** Length of all SH-SY5Y and sequin pass reads. Dashed line shows median read length. X-axis truncated at 5 kb. **(C)** Gene body coverage of SH-SY5Y reads in each sequenced sample. Length of all genes normalised to 100. 5’ end of gene on left-hand side (0) to 3’ end on right-hand side (100). Lines show mean coverage across all genes across the length of the gene body. U1 & U2 - Undifferentiated replicates 1 & 2. D1 & D3 - Differentiated replicates 1 & 3. D2T1 & T2 - Technical replicates of differentiated replicate sample 2. **(D)** Fraction of GENCODE v31 transcripts covered by each read compared to annotated transcript length. Density plot shows the annotated length and the transcript fraction covered by each SH-SY5Y read. The trend line of coverage vs transcript length was plotted using a generalized additive model, an extension of a generalised linear model where the linear form is replaced by sum of smooth functions. **(E)** Fraction of full-length SH-SY5Y reads. Fraction of GENCODE v31 transcript covered by each read. Dotted line represents 95% cutoff for full-length reads. Full-length reads shaded in blue. X-axis truncated at 50% coverage.

### Sequencing metrics and read alignment

dRNA sequencing generated 6.5 million reads in total, of which 4.4 million (∼68%) were pass reads with a quality score greater than 7. Yeast calibration RNA generated 325k pass reads which had a median read length of 1.3 kb, which was consistent with the known length of this control and a median accuracy of 91%. There were ∼4.1 million pass reads from 5Y cells and RNA sequins (hereafter referred to as “reads”). Reads had a median length of 1004 nucleotides and a median quality score of 10 across all samples (**Figure 1B, Table 1**).

**Table 1.** dRNA sequencing and alignment metrics (pass reads only)

|  | Undiff1 | Undiff2 | Diff1 | Diff2TR1 | Diff2TR2 | Diff3 | Overall |
| --- | --- | --- | --- | --- | --- | --- | --- |
| Number of reads | 468,790 | 778,037 | 518,176 | 806,067 | 867,371 | 634,226 | 4,072,667 |
| Median read length (nt) | 989 | 1,009 | 922 | 994 | 1,041 | 1,042 | 1,004 |
| Median read quality score | 9.8 | 10.2 | 9.9 | 10.0 | 10.0 | 9.9 | 10.0 |
| Longest alignment length (nt) | 13,439 | 15,081 | 12,600 | 13,739 | 13,624 | 14,362 | 15,081 |
| Reads aligned to genome (%) | 97.5 | 97.8 | 97.8 | 98.1 | 98.2 | 98.1 | 98.0 |
| Reads aligned to transcriptome (%) | 97.7 | 97.8 | 98.0 | 98.3 | 98.4 | 98.3 | 98.2 |

Reads from each sample were aligned using minimap2 (Li 2018) to the human and sequin reference genome and transcriptome. The aligned lengths had an overall median of 991 nucleotides (**Figure S1A**), demonstrating almost the entire length of the reads aligned. Ninety eight percent of reads aligned to the genome and transcriptome, meaning almost all reads were assigned to a gene and transcript isoform (**Table 1**). Most reads only had a single primary genomic alignment. Similar to previous findings (Soneson et al. 2019), >70% of reads also had 1 or more secondary transcriptomic alignments, largely reflecting the increased difficulty of assigning reads to specific isoforms. Although isoform assignment was significantly improved by filtering with NanoCount prior to quantification (**Figure S1B**). We identified expression of 17,263 human genes (65% of protein coding genes) and 56,460 human isoforms (42% of protein coding isoforms), as well as 42 (55%) and 78 (49%) sequin genes and isoforms.

### Genome and transcriptome coverage

One of the main advantages of nanopore sequencing is the potential to generate long reads that comprise full-length transcripts. However, both RNA degradation and ONT software limitations reduce the extent to which dRNA reads represent full-length transcripts (Workman et al. 2019). Sequin RNAs are in-vitro transcribed and so are not vulnerable to degradation by cellular RNases during RNA extraction. As such they should provide some indication of how much degradation is present in cellular RNA (even when RNA integrity appears high) vs how many incomplete transcripts are due to degradation during library preparation and/or sequencing limitations.

We examined read coverage across gene bodies. dRNA sequencing requires an intact 3’ end, which is sequenced first (Garalde et al. 2018) and we observed corresponding high coverage at the 3’ end (**Figure 1C**). Gene body coverage then progressively decreased towards the 5’ end, consistent with previous studies (Depledge et al. 2019; Soneson et al. 2019).

To assess the fraction of transcripts covered by reads and the proportion that represent full-length transcripts, we created a dataset of known/annotated transcript lengths from GENCODE (v31) (Frankish et al. 2018). A ‘coverage fraction’ was calculated for each primary alignment to the transcriptome. We defined coverage fraction as the observed transcript length divided by the known transcript length. The median coverage fraction was 0.74 for 5Y RNA, demonstrating that most reads covered a high proportion of the original RNA. This was lower than observed in the sequin data (0.88), suggesting some RNA degradation took place during RNA extraction. Alignments to longer transcripts were less likely to be full-length, consistent with previous studies (Jenjaroenpun et al. 2018) (**Figure 1D, S2B, C**). However, the highest density of coverage was close to 1 for transcripts approximately 1 kb long, showing many shorter transcripts are generally covered in their entirety.

We classified alignments that covered at least 95% of their assigned transcript isoform as full-length as per Jenjaroenpun et al. (2018). In total, 46% of sequin reads were full-length compared to 29% of 5Y reads (**Figure 1E**). These results suggest that while a significant proportion of reads are full-length, there is considerable room for improvement through sequencing software improvements and by preventing degradation during RNA extraction and library preparation.

### Accurate expression quantification with dRNA sequencing

Previous studies using dRNA sequencing revealed good quantitative accuracy (Jenjaroenpun et al. 2018; Garalde et al. 2018; Soneson et al. 2019). To confirm and extend these findings we utilised the sequin controls to investigate the quantitative accuracy of dRNA sequencing, and subsequently the potential to detect differential expression.

Genes and isoforms were quantified with featureCounts (Liao et al. 2013) and NanoCount (Leger 2020) respectively (**Table S1**). To test the quantification accuracy of dRNA sequencing, the observed sequin gene and isoform counts were compared with known sequin input concentrations (**Figure 2A, B**). Measured sequin abundance was highly correlated with known abundance at the gene and isoform level (Spearman’s correlation of 0.96 and 0.91 respectively, both p<0.0001, two-tailed). Results showed dRNA could detect but not quantitate sequins at very low concentrations, therefore a segmental linear regression was utilised to help identify the concentration where quantitative measurement began. This revealed a slope close to 1 above the cutoff for genes and isoforms. These findings demonstrate the ability of dRNA sequencing to accurately quantify detected genes and isoforms.

**Figure 2.**
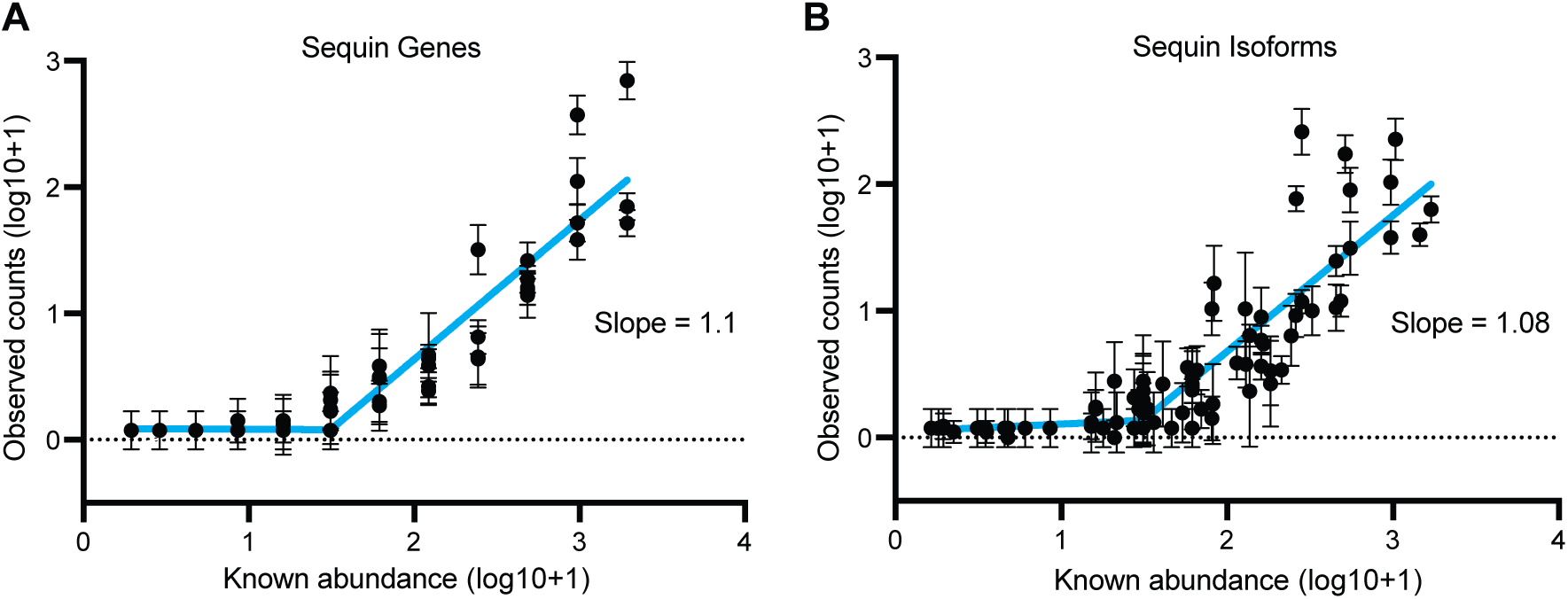
Quantification of sequin spike-in controls. **(A)** Quantification of sequin genes. **(B)** Quantification of sequin transcript isoforms. Comparison of known sequin Mix B abundance (original concentration in attomoles/ul) to measured counts. Concentrations transformed log_10_(X+1). Mean and standard deviation plotted. N = 4. Trend line = segmental linear regression with breakpoint at 1.49 performed on log_10_ transformed data.

### Identification of differential expression with dRNA sequencing

We next asked if dRNA sequencing could detect systematic expression level changes between conditions. Comparison of Mix A and B sequins using principal component analyses (PCA) demonstrated sequin mixes separated on PC1 (**Figure 3A, S3**). Expression changes between the mixes explained a very high degree of the variance (83% at the gene level and 79% at the transcript isoform level). Similarly, endogenous 5Y samples separated by differentiation state along PC1, which explained 82% of the variance at the gene and 76% at the isoform level (**Figure 3B**). Technical replication produced almost identical samples, further confirming little technical variability from library preparation and sequencing (**Figure S3**). Together the sequin and 5Y results illustrate that dRNA sequencing can robustly identify differences in expression between samples at both the gene and isoform level and that the measured expression changes are reflective of the biological changes between samples.

**Figure 3.**
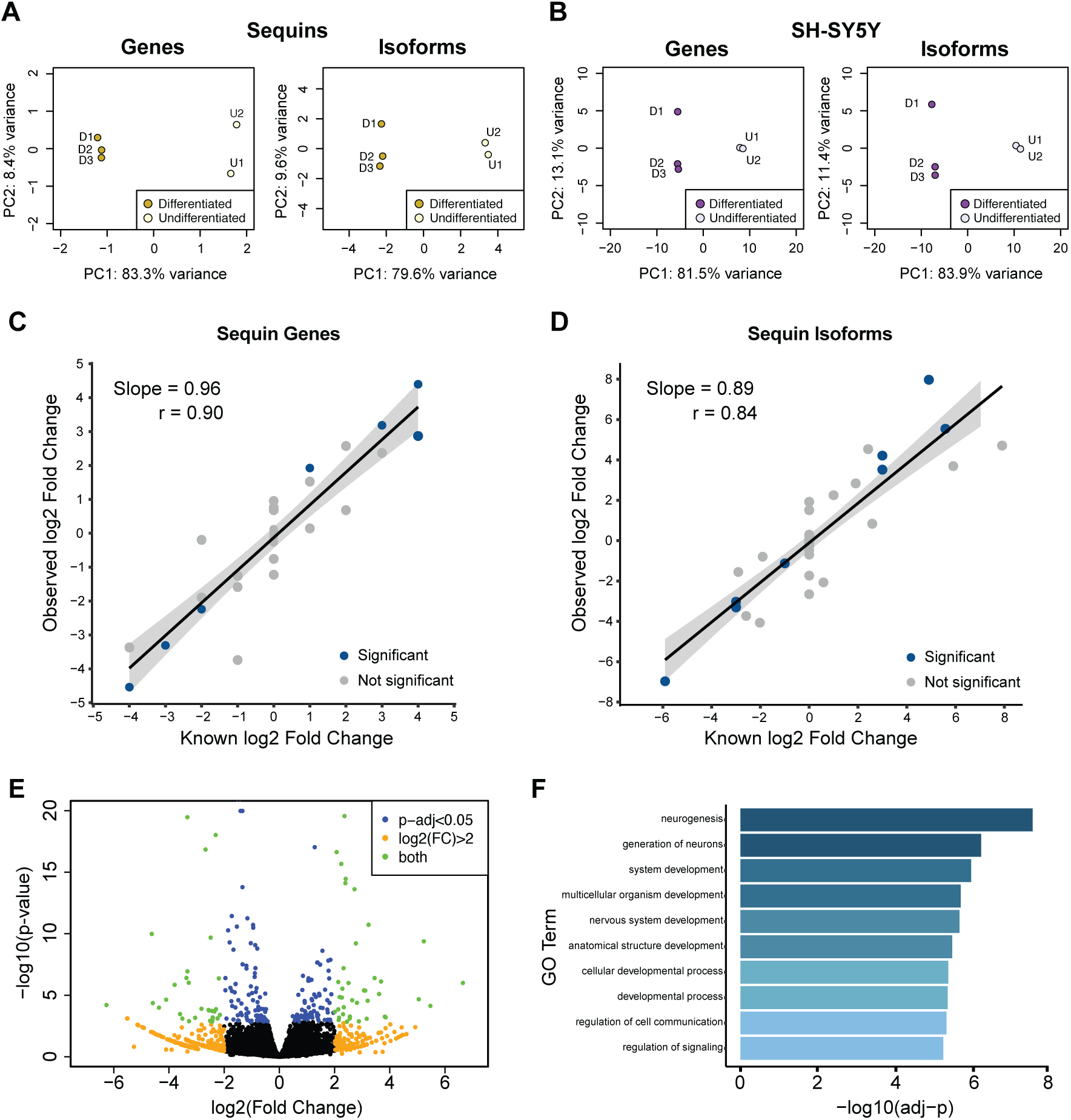
Identification of differential gene and isoform expression. **(A/B)** Principal component analysis (PCA) of sequin (A) and SH-SY5Y (B) gene and isoform expression between undifferentiated and differentiated SH-SY5Y cells. All plots show the first two principal components. SH-SY5Y shows endogenous expression only. Sequins were added to undifferentiated (Mix A) and differentiated (Mix B) SH-SY5Y RNA and plots reflect measured abundance differences between the sequin mixes. **(C/D)** Quantification of fold changes between Mix A and Mix B sequin genes (C) and isoforms (D). Comparison of known fold changes in abundance with measured fold changes from sequencing. Sequins with significant differential expression in blue, not significant in grey. Trend line shows slope from linear regression. Shaded grey area shows 95% confidence interval for regression slope. Correlation (r) is spearman correlation. **(E)** Volcano plot of differential gene expression between undifferentiated and differentiated SH-SY5Y cells. An adjusted p-value<0.05 (from DESeq2) was considered significant for differential expression. FC = Fold Change. **(F)** Gene ontology (GO) terms most associated with differentially expressed genes upregulated during SH-SY5Y differentiation. P-values adjusted using Bonferroni correction for multiple testing.

We examined differential expression (DE) of genes and transcript isoforms using DESeq2 (Love et al. 2014). We identified 7 sequin genes and 8 sequin isoforms as differentially expressed (adj-p<0.05). All were true positives suggesting high accuracy in the results (**Figure 3C,D, Table S2**). There are 47 sequin genes and 83 sequin transcripts potentially detectable as differentially expressed (fold change ≠ 0), however only 18 / 22 of these genes and transcripts passed our filtering thresholds for analysis. This suggests the high specificity, but lower sensitivity may be due to low overall read counts. To examine if dRNA sequencing accurately identified changes in sequin concentrations between Mix A and B, even if significance was not reached, we examined the correlation between observed and known log2 fold changes of sequin genes and isoforms and also performed a linear regression to identify the quantification accuracy of fold change measurement (**Figure 3C,D**). Correlations between observed and known log2 fold changes were high, 0.90 for genes and 0.84 for isoforms, and the slopes were close to one, 0.96 for genes and 0.89 for isoforms (both significant at p<1×10^−12^). The significant linear relationships and strong correlations at both the sequin gene and isoform level demonstrates dRNA sequencing accurately detected changes in expression between groups. The slightly lower values for transcripts isoforms may be due to the increased difficulty in unambiguously assigning a read to a specific isoform, as most sequins, (like human genes) have multiple isoforms **(Figure S1B**).

Applying the DE analysis on 5Y samples identified 231 annotated genes and 291 transcript isoforms as DE between undifferentiated and differentiated samples (adj-p<0.05) (**Figure 3E, Figure S4, Table S3**). Genes significantly upregulated in differentiated samples (n=118) were used in a gene ontology analysis using PANTHER (Thomas et al. 2003). The most significant GO terms were predominantly associated with neuronal development and developmental processes more generally, with the GO term ‘neurogenesis’ (GO:0022008) the most associated (adj-p=2.9×10^−8^) with differentiated 5Y cells (**Figure 3F, Table S4**). GO analysis for upregulated isoforms (n=181) gave similar results, while ranking significant GO terms by fold enrichment also emphasised a number of terms relating to neuron differentiation, neuron projection and axonal development (**Table S4**). These results validate the ability of dRNA sequencing to identify biologically relevant changes in gene expression.

### Differential transcript usage (DTU) analysis between samples

Along with DE of genes (DGE) and transcript isoforms (DTE), changes in isoform usage (DTU) can also be physiologically relevant and reveal important distinctions between cell types and in disease (Scotti and Swanson 2015). DTU involves isoform switches or changes in isoform proportions and can occur even when there is no change in expression at the gene level. As the results of DTE often largely reflect those of DGE, complexity at the isoform level can be masked (Yi et al. 2018). Our results demonstrated dRNA can perform quantitative sequencing of full-length isoforms and hence should be well suited to revealing this complexity.

Transcript counts were used to identify isoform switching using DEXSeq within the IsoformSwitchAnalyzeR package (Vitting-Seerup and Sandelin 2017). After filtering out lowly expressed genes and isoforms and isoform fraction changes of less than 0.1, 4,823 5Y isoforms and 18 sequin isoforms were included for analysis.

We identified 27 endogenous genes and 1 sequin gene (adj-p<0.05) with isoform switching between undifferentiated and differentiated 5Y cells (**Table S5**). The sequin (adj-p=3×10^−13^) was a true positive for DTU, with a similar observed difference in isoform fractions (0.95) to the known difference (0.88) (**Figure 4A**). Less than 20% of genes with DTU showed DGE, while only 40% of DTU isoforms showed DTE, confirming how complexity at the isoform level can be masked and the importance of separately identifying DTU. To identify the potential consequences of isoform switching, the open reading frames, coding potential and protein domains were predicted for each isoform. Consequences ranged from changes to non-coding 5’ transcriptional start sites and 3’UTRs to alterations in coding regions and protein domains or switches between coding and non-coding isoforms.

**Figure 4.**
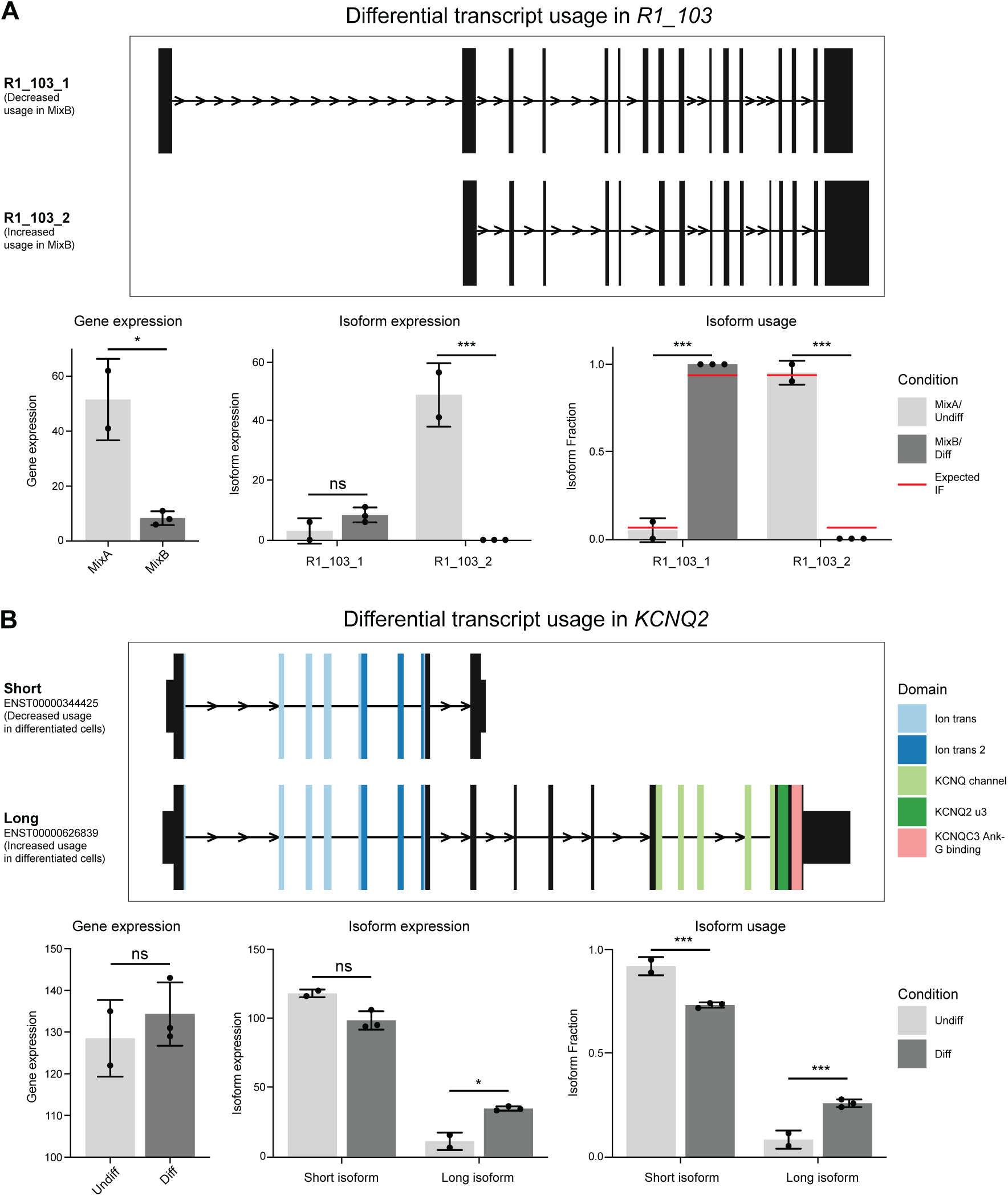
Differential transcript usage of control and endogenous genes. Identification of DTU in the sequin gene *R1_103* **(A)** and the potassium channel KCNQ2 **(B)**. Top: structure of expressed isoforms. Arrows show direction of transcription. Narrow lines show intronic regions (not to scale). Exons displayed as boxes. Taller exonic boxes are coding regions, shorter boxes are 5’ and 3’ UTR regions. Colours represent identified protein domains. Below: graphs display gene and isoform expression and the fraction of expression corresponding to each isoform in Mix A / Undifferentiated and Mix B / Differentiated samples. Red lines show known (expected) sequin isoform fractions in Mix A and B. ns = not significant, * < 0.05, ** < 0.01, *** < 0.001.

DTU identified in endogenous 5Y genes included examples which matched the known biology and expression changes in neuroblastoma cells. *KCNQ2* forms a multimeric potassium channel and expresses multiple mRNA isoforms encoding functionally variant proteins. The short isoform ENST00000344425 (protein isoform 6) lacks much of the cytoplasmic C-terminal region and can alter channel properties to suppress the potassium current (Smith et al. 2001). It is the dominant isoform in undifferentiated IMR-32 neuroblastoma cells and also expressed in developing brain. In contrast, the long isoform ENST00000626839 (protein isoform 2), is the major protein isoform in adult brain and up-regulated in differentiated IMR-32 cells (Smith et al. 2001). While we find no overall change in the expression level of *KCNQ2* in 5Y cells, we identified an expression shift from the short to the long isoform upon differentiation, as would be expected from previous findings (**Figure 4B**). Together these results demonstrate long-read dRNA sequencing can identify and quantify biologically relevant changes in isoform usage.

### Novel transcript identification and analysis

A key advantage of long-read sequencing is its ability to resolve complex RNA isoforms, including the discovery of novel transcripts (Roach et al. 2020; Workman et al. 2019). Due to the ∼10% error rate in dRNA sequencing, confident identification of isoforms first requires correction and clustering of individual reads. We used FLAIR (Tang et al. 2020) to process our aligned reads into a high-confidence set (each represented by at least 5 reads) of 30,914 unique human transcript isoforms (**Figure 5A, Table S6**).

**Figure 5.**
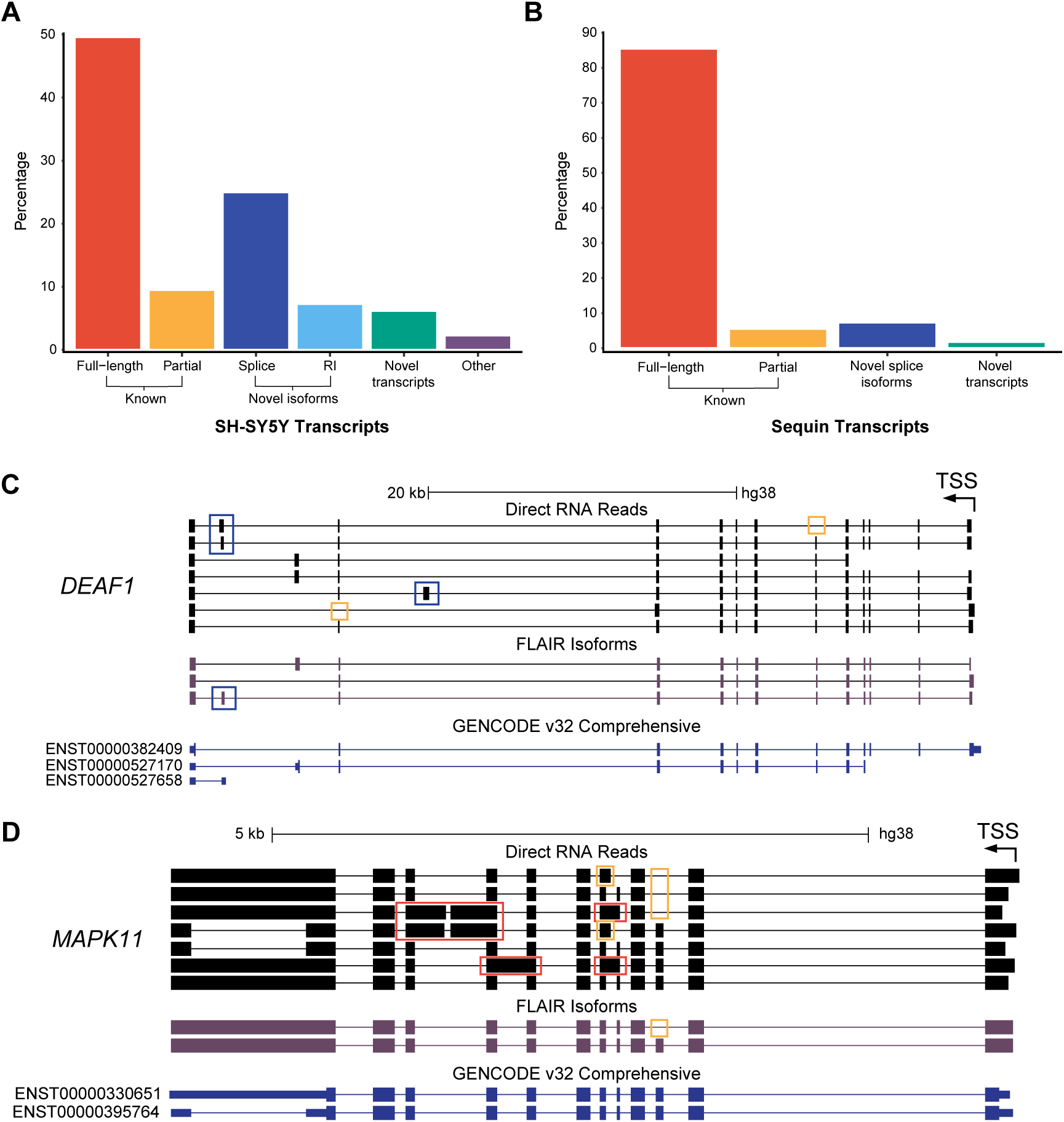
FLAIR transcriptome annotation and identification of novel genes and isoforms. **(A/B)** FLAIR annotation of (A) SH-SY5Y and (B) sequins were compared with known annotations using gffcompare (Pertea and Pertea 2020). The gffcompare class codes were grouped into the following four categories: **known**, transcripts that are full-length or partial-length exact matches to existing annotations; **novel isoform**, transcripts from known genes with novel exon junctions or retained introns (RI); **novel transcript**, novel intergenic, antisense or intronic transcripts potentially representing novel genes; **other**, possible artifacts and RNA fragments. **(C/D)** UCSC Genome Browser screenshots of novel isoforms identified by FLAIR compared to known GENCODE annotations and raw nanopore dRNA reads. *DEAF1* (C) and *MAPK11* (D). Novel exons coloured blue; novel splice junctions of known exons coloured orange; retained intron exons coloured red.

High-confidence FLAIR isoforms were compared to reference annotations using gffcompare (Pertea and Pertea 2020). Isoforms were grouped into the following categories; known, novel isoform (of a known gene), novel gene/transcript and other (including sequencing artefacts). The most abundant category (59.1%) was known isoforms, comprised of full-length matches (49.6%) and partial matches (9.5%) to annotated isoforms. Novel isoforms of known genes (32.3%) largely consisted of novel splice isoforms as opposed to transcripts with retained introns. Although long intronic-retaining transcripts may have been more prone to degradation making their detection less likely. Novel intergenic, intronic or antisense loci accounted for 6.2% of transcripts. Overall, the long-read nature of dRNA sequencing allowed us to identify thousands of novel gene isoforms and potentially novel transcribed loci. Previous long-read studies using dRNA sequencing with FLAIR observed higher percentages of novel isoforms (∼50%) but used a lower read threshold for identification (Soneson et al. 2019; Workman et al. 2019).

Given the high error rate of dRNA sequencing, we used the sequins to estimate what proportion of novel isoforms identified with FLAIR were false positives (**Figure 5B**). All sequin isoforms are annotated and should therefore be classed as known, whereas any novel isoforms represent false positives. FLAIR identified 55 unique sequin isoforms, 50 (91%) known; 4 (7%) novel splice isoforms and 1 (2%) novel intronic transcript. The main source of error was the incorrect identification of exon skipping creating spurious splice isoforms. Additionally, applying the less stringent isoform identification settings used in previous studies increased the false positive rate from 9% to 14%. Taken together these results suggest most novel 5Y isoforms are real but that the false positive rate for novel transcripts classified by FLAIR/gffcompare is quite high.

We investigated examples of genes identified as having novel isoforms with FLAIR (**Figure 5C,D**). *DEAF1* is a transcription factor essential for nervous system development (Hahm et al. 2004). Mutations in *DEAF1* lead to intellectual disability (Vulto-van Silfhout et al. 2014). In agreement with the individual dRNA reads, FLAIR identified the canonical isoform (ENST00000382409) and also provided evidence that two other GENCODE transcripts are likely incomplete isoforms using the standard 5’ initiation site. Other potential novel isoforms were filtered out by FLAIR due to a lack of supporting reads (**Figure 4C**). *MAPK11*, is a protein kinase involved in balancing apoptosis and autophagy (Beardmore et al. 2005). Individual reads suggested a number of novel isoforms including several with retained introns. FLAIR identified the canonical isoform and a novel isoform with skipping of exon 3 (**Figure 5D**).

## Discussion

To enable wide uptake in the transcriptomics field, dRNA sequencing needs to quantify RNA expression accurately and identify differential expression between tissues, development stages and disease states. We performed an in-depth analysis of dRNA sequencing using synthetic spike-in RNAs and human SH-SY5Y (5Y) neuroblastoma cells. We assessed the capabilities of dRNA sequencing including quantification accuracy, differential expression analysis between groups and novel transcript identification. We demonstrated that dRNA sequencing is able to robustly identify differential expression between synthetic RNA mixes and human cell types and produce biologically relevant information. Combined with its previously demonstrated capabilities, the detection of differential gene and isoform expression, as well as switches in isoform usage (DTU), confirms the widespread potential for dRNA in transcriptome profiling.

Previous studies have demonstrated dRNA sequencing is a highly accurate method for quantifying expression of spike-in controls (Garalde et al. 2018; Sessegolo et al. 2019) and could identify differential gene expression in yeast (Jenjaroenpun et al. 2018). DGE and/or DTE have recently been reported in *C*.*elegans, Arabidopsis* and mammalian cells with dRNA (Zhang et al. 2020; Li et al. 2020; Robinson et al. 2020). However, these have almost exclusively been identified from fold changes alone or using very lenient statistics, leaving the capability of dRNA to identify differential expression in a robust manner uncertain. In addition, the human transcriptome (with over 200,000 annotated transcript isoforms) is vastly more complex than that of model organisms (Frankish et al. 2018). This creates a larger challenge both in terms of sequencing depth required to quantify human genes and isoforms but also in performing accurate assignment of reads to their correct transcript isoform. Despite the modest read depth obtained, we found dRNA was sensitive enough to detect significant expression differences in 231 genes and 291 isoforms along with 27 isoform switches during 5Y differentiation. In both human and sequin data, samples clustered by group in PC1 at both the gene and isoform level. The subsequent GO analysis showed that DE genes in differentiated 5Y cells were associated with neurogenesis. We used our sequin control to validate a close relationship between known and measured fold changes even if the significance threshold for DE wasn’t reached. Hence, we expect that the majority of 5Y results represent true positives, but that the results are likely missing a number of DE genes and isoforms that may be of biological importance.

Although the number of replicates used in this analysis was lower than recommended (Schurch et al. 2016), the technique still identified many differentially expressed features between groups. This raises the possibility that the number of replicates required to identify differential expression may be lower for dRNA sequencing due to the low level of bias and possibility of highly accurate quantification.

The key challenges with dRNA are sequencing depth and correct isoform assignment. The first is due to the modest number of reads generated per sequencing run (0.5-1 million). Although higher sequencing depths with >1 million pass reads per run are now possible, it remains challenging for dRNA sequencing to detect lowly expressed genes and isoforms. This is demonstrated in the sequin results and indicates that many 5Y genes and isoforms are below the level for quantitative detection. The read depth achievable with dRNA sequencing needs to increase to routinely out-perform short-read RNA-seq in sensitivity. Currently, the ideal method for comprehensive transcriptome analysis is to complement high-throughput short-read sequencing with long-read technologies (Weirather et al. 2017).

The challenge of correct isoform assignment has been previously noted (Sessegolo et al. 2019; Soneson et al. 2019) and is due to the high (∼10%) error rate and the large proportion of non-full-length reads. Although RNA extracted from 5Y cells was of very high quality (RIN>9), it remains challenging to sequence a high proportion of full-length transcripts. dRNA sequencing starts at the 3’ end of transcripts (Garalde et al. 2018), therefore many incomplete reads are largely consigned to 3’UTRs and cannot specify the expressed isoform. Exemplifying this, >70% of reads had secondary transcriptome alignments, although quantification programs such as NanoCount use filtering and expectation–maximization to improve isoform assignment. For this reason, we do not recommend using primary alignments alone as this causes many reads to be specifically assigned to incorrect isoforms. Methodological improvements to increase the proportion of full-length transcripts will be just as important to maximising correct transcript assignment, if not more so, than those which improve dRNA sequencing accuracy. Another recent dRNA study reported that approximately 20% of reads are truncated during sequencing due to signal noise (Workman et al. 2019). The truncation of reads due to software limitations is an obvious avenue for improvement, along with preventing RNA degradation during library preparation, for maximising full-length transcripts.

A key advantage of dRNA sequencing is its ability to sequence transcripts as they exist in the cell, improving the detection of novel transcript isoforms and RNA modifications. In contrast, Illumina short-read sequencing performs poorly in resolving transcript isoforms, especially those resulting from alternative splicing (Steijger et al. 2013). FLAIR (Tang et al. 2020) identified ∼31,000 isoforms in 5Y cells, 38.5% of which were novel isoforms of known genes or potentially novel transcriptional units. Inclusion of spike-in controls provides confidence that most of these novel transcripts are genuine. The high number of novel transcripts detected in this study and others indicates that the human transcriptome annotation is far from complete. Improving this annotation is a task in which dRNA sequencing will be highly valuable.

In summary, the applicability of dRNA sequencing to gene expression profiling was demonstrated using synthetic controls and human cell populations. We show that dRNA sequencing identifies biologically relevant changes in gene and isoform expression in complex transcriptomes. The long read lengths generated by the technique provide a clear advantage for isoform quantification, however the throughput and accuracy of dRNA sequencing requires improvement before it can truly outweigh other sequencing technologies. In conclusion, dRNA is a promising method to decipher the complex expression and splicing patterns that characterise the transcriptome and to identify differentially expressed isoforms contributing to development and disease.

## Methods

### Cell culture

Human SH-SY5Y neuroblastoma cells were cultured for use as a model of neuronal differentiation (Biedler et al. 1978). Undifferentiated SH-SY5Y cells were cultured in growth media under standard conditions (5% CO2, 37°C) with DMEM:F12 (Sigma D6421), supplemented with 10% fetal bovine serum (FBS) (Sigma F9665), 2 mM L-glutamine and 1% non-essential amino acids. SH-SY5Y differentiations were performed in triplicate. To differentiate the SH-SY5Y cells, flasks were coated with poly-lysine. Cells were seeded at 4.4×10^4^ cells/cm^2^ and grown for 24 hours in standard growth medium. The growth media was then exchanged for differentiation media neurobasal medium (Thermo Fisher 21103049), 2 mM L-glutamine, 1% FBS, 10% B27 supplement (Thermo Fisher 17504044) and 10 mM retinoic acid (Sigma R2625). Media was replaced after 48 hours. After 72 hours of exposure to 10 mM retinoic acid the media was exchanged to differentiation media without retinoic acid and cells were allowed to further differentiate for 72 hours, including a media change at 48 hours. RNA was then extracted from undifferentiated and differentiated cells with Tri Reagent (Thermo Fisher 15596026) and RNeasy columns (Qiagen 74106).

### Library preparation and sequencing

To prepare RNA for sequencing, Turbo DNase was added for 30 minutes at 37°C in a PCR thermocycler with the lid set to 50°C to remove any DNA contamination. Samples were purified with the RNeasy MinElute Cleanup Kit (Qiagen 74204) and eluted with 15 μl nuclease-free water. Samples were run on an Agilent 4200 Tapestation to ensure RNA had a RIN of >9 and quantified via Qubit (Thermo Fisher). PolyA+ purification was performed with 50 μl of NEXTflex polyA+ beads on a minimum of 40 μg of total RNA. Confirmation of rRNA removal and sample quantification were performed with Qubit and Tapestation respectively. To minimise batch effects, all samples were prepared together.

Sequencing libraries were prepared with the SQK-RNA001 kit (Oxford Nanopore Technologies, ONT) using the standard protocol, including the optional reverse transcriptase step. Two different controls were added to each sample, yeast calibration RNA (supplied by ONT) and synthetic sequin V2 spike-in RNAs (Hardwick et al. 2016). The calibration RNA is from the yeast enolase II (YHR174W) gene and is 1.3kb in length. RNA sequins provide a quantitative and qualitative reference to enable transcriptome analysis. Sequins are synthetic spliced RNA transcripts for the investigation of gene and isoform quantification, alternative splicing and differential expression. RNA sequins consist of 160 isoforms from 76 artificial genes that vary in concentration over 4 orders of magnitude. Sequin controls are available in two mixes, Mix A and Mix B, which contain the same synthetic RNA isoforms but at different concentrations. Sequins were added to each sample at a concentration of 6%. Undifferentiated samples (U1, U2) contained sequin Mix A, and differentiated samples (D1, D2TR1, D2TR2, D3) contain Mix B. Sample input was 352.5 ng of polyA+ RNA and 22.5 ng of sequin RNA. Libraries were sequenced on the ONT MinION using R9.4.1 flow cells and MinKNOW (v18.01.6) to generate FAST5 files. FAST5 files were basecalled with Guppy (v3.4.5) (ONT) to create summary text files and FASTQ files of the reads.

### Sequencing metrics and quality control

Initial data analysis was performed to gather general metrics on sequencing performance. All analyses were performed on pass reads (quality (Q) score >= 7). To ensure the sequencing was of high quality, EPI2ME (ONT) was used to determine the sequencing accuracy of the yeast calibration RNA. All subsequent analyses were performed on pass SH-SY5Y and sequin reads (hereafter referred to as reads). Overall data characteristics including median read length; median quality score; longest read length; total number of reads; read length and quality scores over run time were performing using NanoPack (Coster et al. 2018) and pycoQC (Leger and Leonardi 2019).

### Genome and transcriptome alignment

FASTQ files containing ONT pass reads were aligned to the human (GRCh38) (Zerbino et al. 2017) and synthetic (Hardwick et al. 2016) sequin genome and transcriptome using minimap2 (v2.17) (Li 2018). The genome alignments were performed using the splice-aware mode of minimap2 *-ax splice -uf* as recommended. The transcriptome alignments were performed using the long-read mode for ONT data *-ax map-ont -N 100* to retain multiple secondary alignments. When two alignments both have the same scores, minimap2 assigns the primary one at random. In order to maximise the likelihood of identifying the correct isoform of origin, we retained many secondary alignments.

### Full-length transcript identification

A custom script was used to extract data from transcriptome BAM files and identify full-length transcripts. The GENCODE v31 annotation was used as input to assign a known transcript length to each primary alignment (Frankish et al. 2018). Coverage fractions were then calculated by dividing the observed length by the known length for each read’s primary alignment. Reads were required to cover at least 95% of their annotated transcript to be classed as full-length.

### Gene and isoform quantification

FeatureCounts was applied (v1.6.5) (Liao et al. 2013) to the human or sequin genome alignments along with annotations to calculate gene counts with the parameters *-L --primary*. NanoCount (v0.2.3) (Leger 2020) was used for isoform quantification on BAM files of all alignments with the parameters *-p align_score -3 100 --extra_tx_info --discard_supplementary* to obtain counts for human and sequin transcriptome alignments. These NanoCount parameters remove supplementary alignments, alignments with a 3’ end greater than 100 nt away from the annotated 3’ end site and alignments with less than 90% of the best alignment score for that read. Expectation maximisation to identify the most probable isoform was then applied by NanoCount on this filtered set of alignments. Sequin genes and isoforms are present at known concentrations over a 32773 concentration fold range (genes), and a 229409 fold range (Mix B isoforms) and with known fold changes between Mix A and Mix B. To assess the quantitative abilities of dRNA sequencing, Mix B sequin counts at the gene and isoform level were compared to observed counts. Only detected sequins (count>0) were included in this analysis. Segmental linear regression (SLR) was utilised to identify the sequin concentration where quantitative measurement began. Criteria for breakpoint were 1. Multiple sequin genes/isoforms at concentration; 2. >50% detection across all replicates at concentration; 3. Slope 1 (before breakpoint) 95% confidence interval should include 0; 4. Optimal or near optimal SLR goodness of fit. All sequin Mix B genes and isoforms detected in at least one replicate were used to calculate Spearman correlations. To assess accuracy for detecting fold changes, known fold changes between Mix A and Mix B were compared to observed fold changes. Slopes were determined by linear regression.

### Differential expression analysis

Differential expression analysis was performed with DESeq2 (v1.24) (Love et al. 2014). Normalised counts for the two technical replicates, Diff2TR1 and Diff2TR2, were averaged prior to analysis to produce Diff2 so as not to falsely increase the statistical power between groups. The counts from featureCounts and NanoCount were input for gene and isoform level analysis respectively. Count matrices were filtered to remove very lowly expressed features (≤5 in total for each gene/isoform). Counts were normalised for sequencing depth within DESeq2 prior to statistical analysis. Log2 fold changes and adjusted p-values (using the Benjamini-Hochberg method to correct for multiple testing) were calculated for each annotated gene or isoform and used to determine statistical significance. A regularised log transformation was subsequently performed on the normalised counts for visualisation. The PCA and volcano plots were made using the following code: https://gist.github.com/stephenturner/f60c1934405c127f09a6.

Differential transcript usage (DTU) analysis was performed using an R package IsoformSwitchAnalyzeR (Vitting-Seerup and Sandelin 2017). The isoform counts from NanoCount were input, along with the annotation and transcriptome files. The DTU statistical analysis was performed within IsoformSwitchAnalyzeR with DEXSeq (v1.32) (Anders et al. 2012; Ritchie et al. 2015) to identify exons (used to infer transcript isoforms) present in different proportions between groups. Counts were filtered to remove single isoform genes and a gene and isoform expression cutoff of 5 was included to filter out lowly expressed features. A cutoff fraction of 0.1 was used as the minimum difference in isoform fraction between conditions to further increase stringency. DEXSeq normalises counts and outputs a table of isoform fractions and their adjusted p-values for switches. The coding potential and protein domains of transcripts were then predicted as part of IsoformSwitchAnalyzeR using CPAT and PFAM respectively (Wang et al. 2013; Punta et al. 2011). Premature termination codons and nonsense-mediated decay sensitivity were also predicted as part of the workflow which enabled the functional consequences of isoform switches between conditions to be predicted and plots to be produced of isoform switches. Known isoform fractions in sequin data were calculated by dividing each isoform concentration by its total gene concentration. To compare isoform switches with DGE and DTE in the same genes (**Table S5**) we used consistent counts data to calculate adjusted p-values. DTE results from DESeq2 (described above) were input directly, while DGE was performed with DESeq2 after collapsing isoform counts to gene counts with IsoformSwitchAnalyzeR.

### Novel transcript identification

FLAIR (v1.4) was used on the human and sequin primary genome alignments (BAM converted to BED12 files using FLAIR script) for novel transcript identification (Tang et al. 2020). Samples were first corrected using genome annotations and with the *-n* flag enabled to keep read strands consistent after correction. The two separate corrected PSL files for human and sequin data were then collapsed into high-confidence isoforms that were represented by at least 5 reads.

The collapsed isoform GTF files were compared to annotated transcripts using gffcompare (v0.11.2) (Pertea and Pertea 2020). Gffcompare assigns each transcript a class code based on how it compares to annotated reference transcripts. The class codes were grouped into the following categories. Known: full (=) or partial (c) intron chain match to annotated transcript isoform. Novel isoforms of known gene: novel splice isoform (j), splice chain match with additional terminal exon(s) (k), retained intron isoforms (m,n). Novel transcripts potentially representing novel genes or transcriptional units: intronic transcript (i), antisense transcript (x), intergenic transcript (u), 5’ distal overlapping RNA (y), other exonic overlap (o). Other: including RNA fragments and potential sequencing artifacts (p,e,s,r).

## Data and code availability

All data, including FAST5 and FASTQ outputs is available from ENA: PRJEB39347.

All code used to perform the analysis (custom scripts, R workflows for DGE, DTE and DTU) is available on GitHub: https://github.com/josiegleeson/directRNA; https://github.com/josiegleeson/BamSlam.

## Acknowledgments

The authors would like to thank Dr Ricardo de Paoli-Iseppi and other members of the Clark lab for useful feedback on the manuscript; Jarny Choi from the Centre for Stem Cell Systems and the Ritchie lab (WEHI) for advice and assistance with analysis methodologies.

## Author Contributions

M.B.C conceived the study. M.B.C, P.H and W.H designed experiments. T.L performed cell culture. M.B.C prepared RNA for sequencing and performed nanopore sequencing. J.G performed bioinformatic analysis with assistance from W.H and M.B.C. M.B.C and J.G wrote the manuscript with assistance from all authors.

## References

Aird D, Ross MG, Chen W-S, Danielsson M, Fennell T, Russ C, Jaffe DB, Nusbaum C, Gnirke A. 2011. Analyzing and minimizing PCR amplification bias in Illumina sequencing libraries. Genome Biology 12: R18.

Anders S, Reyes A, Huber W. 2012. Detecting differential usage of exons from RNA-seq data. Genome Research 22: 2008–2017.

Beardmore VA, Hinton HJ, Eftychi C, Apostolaki M, Armaka M, Darragh J, McIlrath J, Carr JM, Armit LJ, Clacher C, et al. 2005. Generation and Characterization of p38β (MAPK11) Gene-Targeted Mice. Molecular and Cellular Biology 25: 10454–10464.

Biedler JL, Roffler-Tarlov S, Schachner M, Freedman LS. 1978. Multiple neurotransmitter synthesis by human neuroblastoma cell lines and clones. Cancer research 38: 3751–7.

Cali DS, Kim JS, Ghose S, Alkan C, Mutlu O. 2019. Nanopore Sequencing Technology and Tools for Genome Assembly: Computational Analysis of the Current State, Bottlenecks and Future Directions. Briefings in bioinformatics 4: 1542–1559.

Coster W de, D’Hert S, Schultz DT, Cruts M, Broeckhoven C van. 2018. NanoPack: visualizing and processing long-read sequencing data. Bioinformatics 34: 2666–2669.

Depledge DP, Srinivas KP, Sadaoka T, Bready D, Mori Y, Placantonakis DG, Mohr I, Wilson AC. 2019. Direct RNA sequencing on nanopore arrays redefines the transcriptional complexity of a viral pathogen. Nature Communications 10: 754.

Emilsson V, Thorleifsson G, Zhang B, Leonardson AS, Zink F, Zhu J, Carlson S, Helgason A, Walters GB, Gunnarsdottir S, et al. 2008. Genetics of gene expression and its effect on disease. Nature 452: 423–8.

Frankish A, Diekhans M, Ferreira A-M, Johnson R, Jungreis I, Loveland J, Mudge JM, Sisu C, Wright J, Armstrong J, et al. 2018. GENCODE reference annotation for the human and mouse genomes. Nucleic Acids Research 47: D766–D773.

Garalde DR, Snell EA, Jachimowicz D, Sipos B, Lloyd JH, Bruce M, Pantic N, Admassu T, James P, Warland A, et al. 2018. Highly parallel direct RNA sequencing on an array of nanopores. Nature Methods 15: 201–206.

Hahm K, Sum EYM, Fujiwara Y, Lindeman GJ, Visvader JE, Orkin SH. 2004. Defective Neural Tube Closure and Anteroposterior Patterning in Mice Lacking the LIM Protein LMO4 or Its Interacting Partner Deaf-1. Molecular and Cellular Biology 24: 2074–2082.

Hardwick SA, Chen WY, Wong T, Deveson IW, Blackburn J, Andersen SB, Nielsen LK, Mattick JS, Mercer TR. 2016. Spliced synthetic genes as internal controls in RNA sequencing experiments. Nature methods 13: 792–8.

Jenjaroenpun P, Wongsurawat T, Pereira R, Patumcharoenpol P, Ussery DW, Nielsen J, Nookaew I. 2018. Complete genomic and transcriptional landscape analysis using third-generation sequencing: a case study of Saccharomyces cerevisiae CEN.PK113-7D. Nucleic Acids Research 46: 38.

Lee TI, Young RA. 2013. Transcriptional regulation and its misregulation in disease. Cell 152: 1237–51.

Leger A. 2020. NanoCount. https://zenodo.org/badge/latestdoi/142873004.

Leger A, Leonardi T. 2019. pycoQC, interactive quality control for Oxford Nanopore Sequencing. Journal of Open Source Software 4: 1236.

Li H. 2018. Minimap2: pairwise alignment for nucleotide sequences. Bioinformatics 34: 3094–3100.

Li R, Ren X, Ding Q, Bi Y, Xie D, Zhao Z. 2020. Direct full-length RNA sequencing reveals unexpected transcriptome complexity during Caenorhabditis elegans development. Genome Research 30.

Liao Y, Smyth GK, Shi W. 2013. featureCounts: an efficient general purpose program for assigning sequence reads to genomic features. Bioinformatics 30: 923–930.

Liu H, Begik O, Lucas MC, Ramirez JM, Mason CE, Wiener D, Schwartz S, Mattick JS, Smith MA, Novoa EM. 2019. Accurate detection of m6A RNA modifications in native RNA sequences. Nature Communications 10: 4079.

Lorenz DA, Sathe S, Einstein JM, Yeo GW. 2019. Direct RNA sequencing enables m 6 A detection in endogenous transcript isoforms at base-specific resolution. RNA 26: 19–28.

Love MI, Huber W, Anders S. 2014. Moderated estimation of fold change and dispersion for RNA-seq data with DESeq2. Genome Biology 15: 550.

Melé M, Ferreira PG, Reverter F, DeLuca DS, Monlong J, Sammeth M, Young TR, Goldmann JM, Pervouchine DD, Sullivan TJ, et al. 2015. The human transcriptome across tissues and individuals. Science 348: 660–665.

Pan Q, Shai O, Lee LJ, Frey BJ, Blencowe BJ. 2008. Deep surveying of alternative splicing complexity in the human transcriptome by high-throughput sequencing. Nature Genetics 40: 1413–1415.

Pertea G, Pertea M. 2020. GFF Utilities: GffRead and GffCompare. F1000Research 9: 304.

Punta M, Coggill PC, Eberhardt RY, Mistry J, Tate J, Boursnell C, Pang N, Forslund K, Ceric G, Clements J, et al. 2011. The Pfam protein families database. Nucleic Acids Research 40: D290–D301.

Ritchie ME, Phipson B, Wu D, Hu Y, Law CW, Shi W, Smyth GK. 2015. limma powers differential expression analyses for RNA-sequencing and microarray studies. Nucleic Acids Research 43: e47–e47.

Roach NP, Sadowski N, Alessi AF, Timp W, Taylor J, Kim JK. 2020. The full-length transcriptome of C. elegans using direct RNA sequencing. Genome Research 30: 299–312.

Robinson EK, Jagannatha P, Covarrubias S, Cattle M, Safavi R, Song R, Viswanathan K, Shapleigh B, Abu-Shumays R, Jain M, et al. 2020. Inflammation Drives Alternative First Exon usage to Regulate Immune Genes including a Novel Iron Regulated Isoform of Aim2. bioRxiv. https://www.biorxiv.org/content/10.1101/2020.07.06.190330v1.

Roundtree IA, He C. 2016. RNA epigenetics—chemical messages for posttranscriptional gene regulation. Current Opinion in Chemical Biology 30: 46–51.

Schurch NJ, Schofield P, Gierliński M, Cole C, Sherstnev A, Singh V, Wrobel N, Gharbi K, Simpson GG, Owen-Hughes T, et al. 2016. How many biological replicates are needed in an RNA-seq experiment and which differential expression tool should you use? RNA 22: 839–851.

Scotti MM, Swanson MS. 2015. RNA mis-splicing in disease. Nature Reviews Genetics 17: 19–32.

Sessegolo C, Cruaud C, Silva C da, Cologne A, Dubarry M, Derrien T, Lacroix V, Aury J-M. 2019. Transcriptome profiling of mouse samples using nanopore sequencing of cDNA and RNA molecules. Scientific Reports 9: 14908.

Sharon D, Tilgner H, Grubert F, Snyder M. 2013. A single-molecule long-read survey of the human transcriptome. Nature Biotechnology 31: 1009–1014.

Smith JS, Iannotti CA, Dargis P, Christian EP, Aiyar J. 2001. Differential Expression of KCNQ2 Splice Variants: Implications to M Current Function during Neuronal Development. Journal of Neuroscience 21: 1096–1103.

Soneson C, Yao Y, Bratus-Neuenschwander A, Patrignani A, Robinson MD, Hussain S. 2019. A comprehensive examination of Nanopore native RNA sequencing for characterization of complex transcriptomes. Nature Communications 10: 3359.

Steijger T, Abril JF, Engström PG, Kokocinski F, Abril JF, Akerman M, Alioto T, Ambrosini G, Antonarakis SE, Behr J, et al. 2013. Assessment of transcript reconstruction methods for RNA-seq. Nature Methods 10: 1177–1184.

Stephenson W, Razaghi R, Busan S, Weeks KM, Timp W, Smibert P. 2020. Direct detection of RNA modifications and structure using single molecule nanopore sequencing. bioRxiv. https://www.biorxiv.org/content/10.1101/2020.05.31.126763v1.

Sui X, Kong N, Ye L, Han W, Zhou J, Zhang Q, He C, Pan H. 2014. p38 and JNK MAPK pathways control the balance of apoptosis and autophagy in response to chemotherapeutic agents. Cancer Letters 344: 174–179.

Tang AD, Soulette CM, Baren MJ van, Hart K, Hrabeta-Robinson E, Wu CJ, Brooks AN. 2020. Full-length transcript characterization of SF3B1 mutation in chronic lymphocytic leukemia reveals downregulation of retained introns. Nature Communications 11: 1438.

Thomas PD, Campbell MJ, Kejariwal A, Mi H, Karlak B, Daverman R, Diemer K, Muruganujan A, Narechania A. 2003. PANTHER: A Library of Protein Families and Subfamilies Indexed by Function. Genome Research 13: 2129–2141.

Vitting-Seerup K, Sandelin A. 2017. The Landscape of Isoform Switches in Human Cancers. Molecular Cancer Research 15: 1206–1220.

Vulto-van Silfhout AT, Rajamanickam S, Jensik PJ, Vergult S, de Rocker N, Newhall KJ, Raghavan R, Reardon SN, Jarrett K, McIntyre T, et al. 2014. Mutations Affecting the SAND Domain of DEAF1 Cause Intellectual Disability with Severe Speech Impairment and Behavioral Problems. The American Journal of Human Genetics 94: 649–661.

Wang ET, Sandberg R, Luo S, Khrebtukova I, Zhang L, Mayr C, Kingsmore SF, Schroth GP, Burge CB. 2008. Alternative isoform regulation in human tissue transcriptomes. Nature 456: 470–476.

Wang L, Park HJ, Dasari S, Wang S, Kocher J-P, Li W. 2013. CPAT: Coding-Potential Assessment Tool using an alignment-free logistic regression model. Nucleic Acids Research 41: e74–e74.

Weirather JL, Cesare M de, Wang Y, Piazza P, Sebastiano V, Wang X-J, Buck D, Au KF. 2017. Comprehensive comparison of Pacific Biosciences and Oxford Nanopore Technologies and their applications to transcriptome analysis. F1000Research 6: 100.

Workman RE, Tang AD, Tang PS, Jain M, Tyson JR, Razaghi R, Zuzarte PC, Gilpatrick T, Payne A, Quick J, et al. 2019. Nanopore native RNA sequencing of a human poly(A) transcriptome. Nature methods 16: 1297–1305.

Yi L, Pimentel H, Bray NL, Pachter L. 2018. Gene-level differential analysis at transcript-level resolution. Genome Biology 19: 53.

Zerbino DR, Achuthan P, Akanni W, Amode MR, Barrell D, Bhai J, Billis K, Cummins C, Gall A, Girón CG, et al. 2017. Ensembl 2018. Nucleic Acids Research 46: D754–D761.

Zhang S, Li R, Zhang L, Chen S, Xie M, Yang L, Xia Y, Foyer CH, Zhao Z, Lam H-M. 2020. New insights into Arabidopsis transcriptome complexity revealed by direct sequencing of native RNAs. Nucleic Acids Research gkaa588.

