## Supplementary_figures for "Nanopore direct RNA sequencing detects differential expression between human cell populations"

#### Figure S1

### A

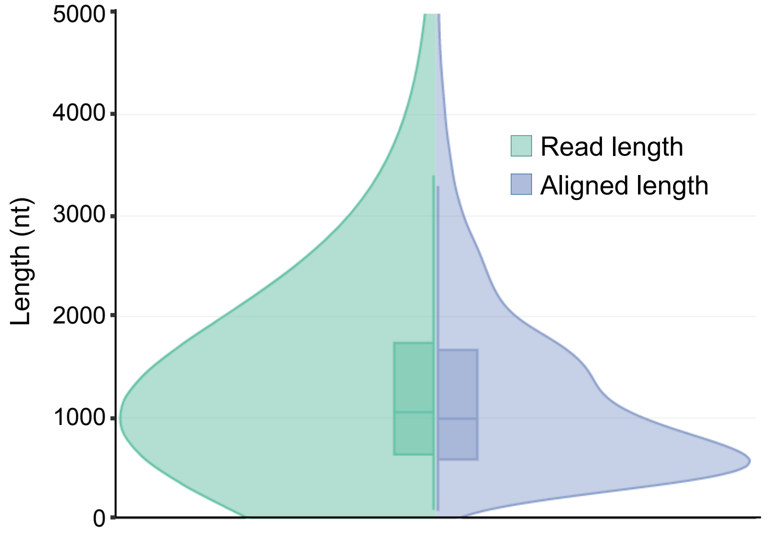


### B

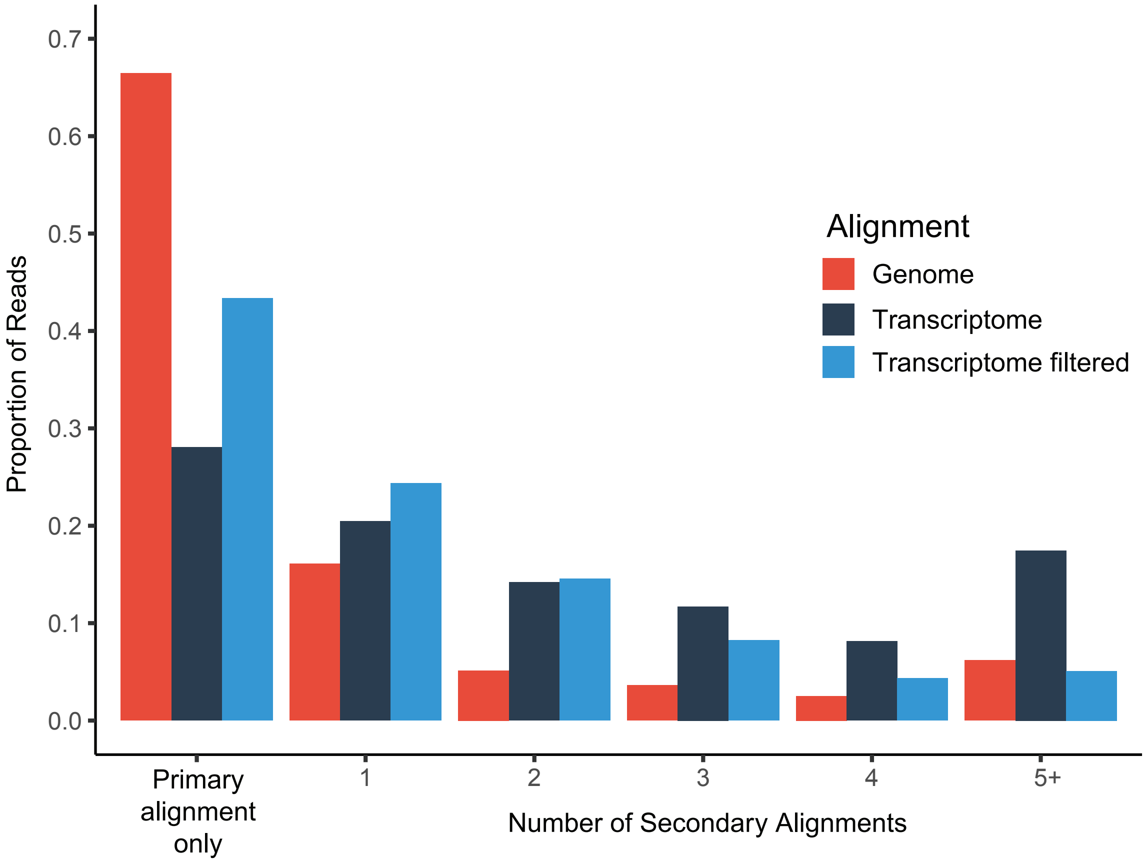


**Figure S1: Alignment of nanopore dRNA reads. (A)** Violin plot split into raw read lengths (green) and the length of the aligned portion of each read to the human transcriptome (purple), for SHSY5Y data. Aligned lengths are plotted for the ~98% of reads that align to the transcriptome. Box plots display length quartiles with density distributions on the outside. **(B)** Proportion of aligned reads that have one or more alignments to the genome and transcriptome. Filtering transcriptome alignments with increased stringency was performed prior to quantification and improves transcript assignment. All plotted reads have a primary alignment and may also have one or more secondary alignments (see methods).

#### Figure S2

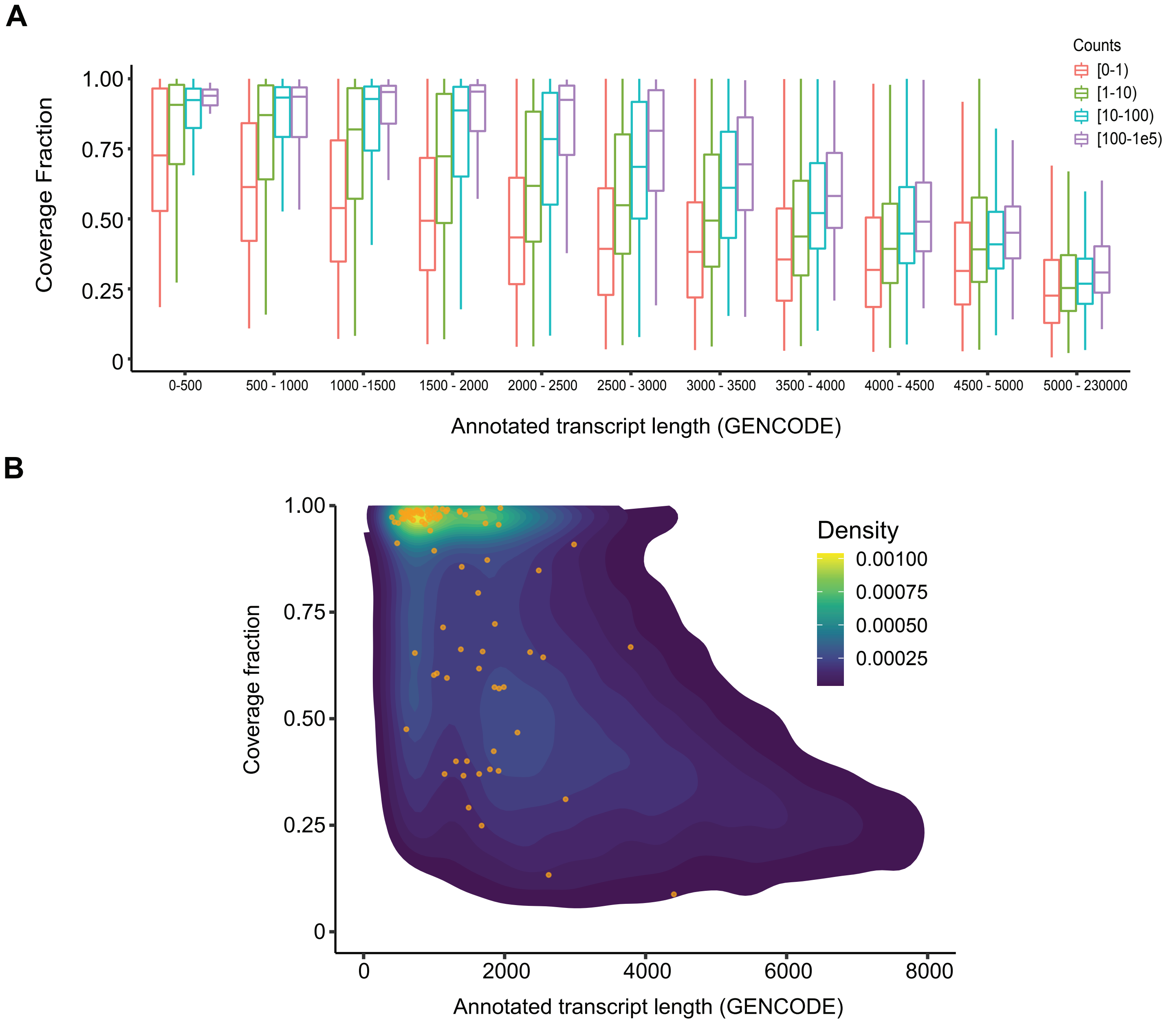


**Figure S2: Transcript coverage vs transcript length. (A)** Fraction of Gencode transcripts covered by aligned reads in different length categories and by number of mapping reads. **(B)** Density plot showing median read coverage of each Gencode transcript compared to annotated transcript length. Orange circles display median coverage fraction for each Sequin transcript. A & B are calculated as the median value per transcript instead of per read (as in Figure 1D).

#### Figure S3

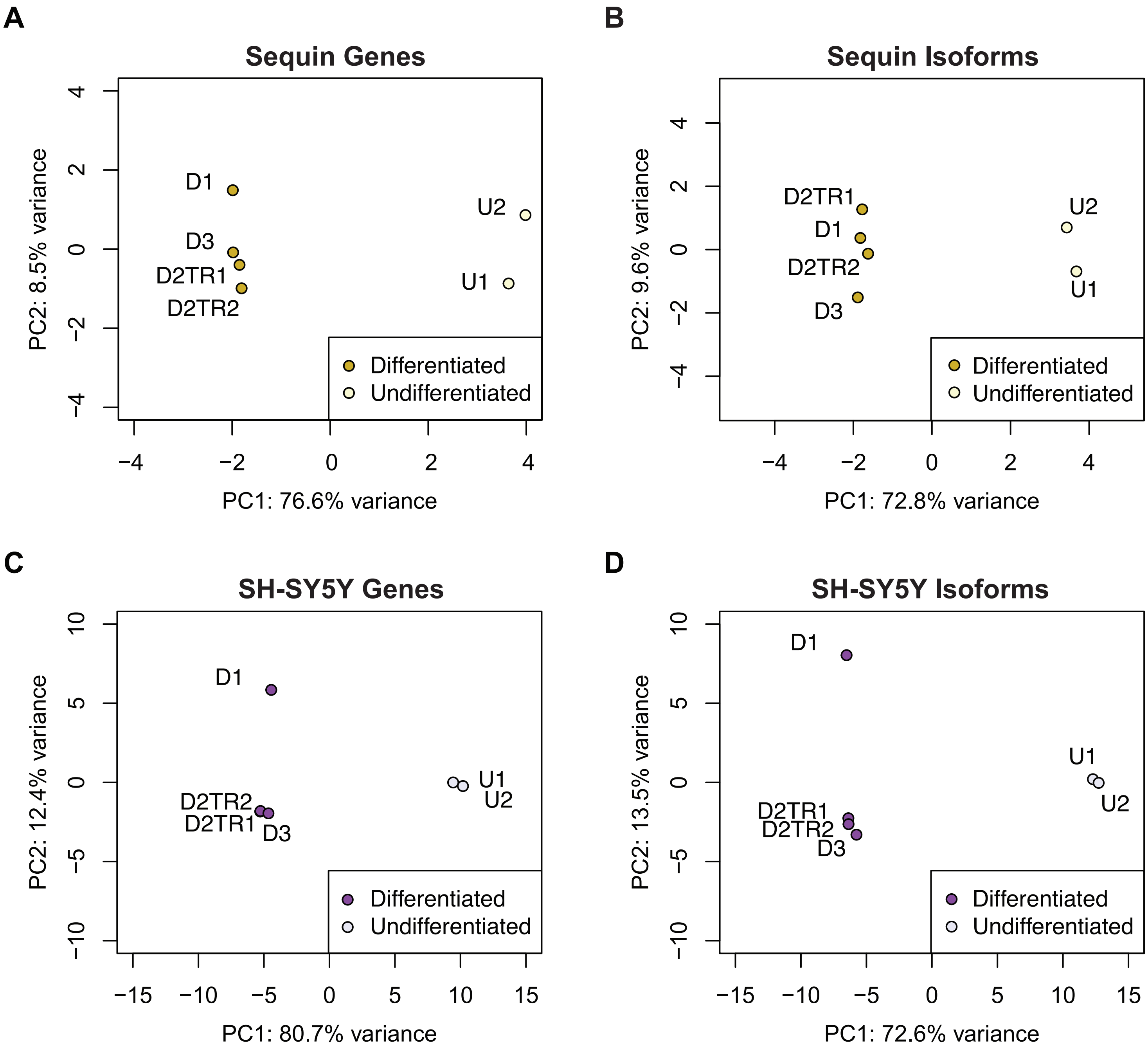


**Figure S3**. Principal component analysis (PCA) of Sequin **(A,B)** and SH-SY5Y **(C,D)** gene and isoform expression, including the technical replicate of differentiation sample 2 (D2). TR = Technical Replicate. All plots show the first two principal components. SH-SY5Y shows endogenous expression only. Sequins were added to undifferentiated (MixA) and differentiated (MixB) SH-SY5Y RNA and plots reflect measured abundance differences between the sequin mixes. PCAs were performed on all expressed features (no expression-based filtering of count matrices). All Sequin samples from the same mix are effectively technical replicates. SH-SY5Y technical replicates use the same polyA+ RNA with an idependant library preparation and sequencing run.

### Figure S4

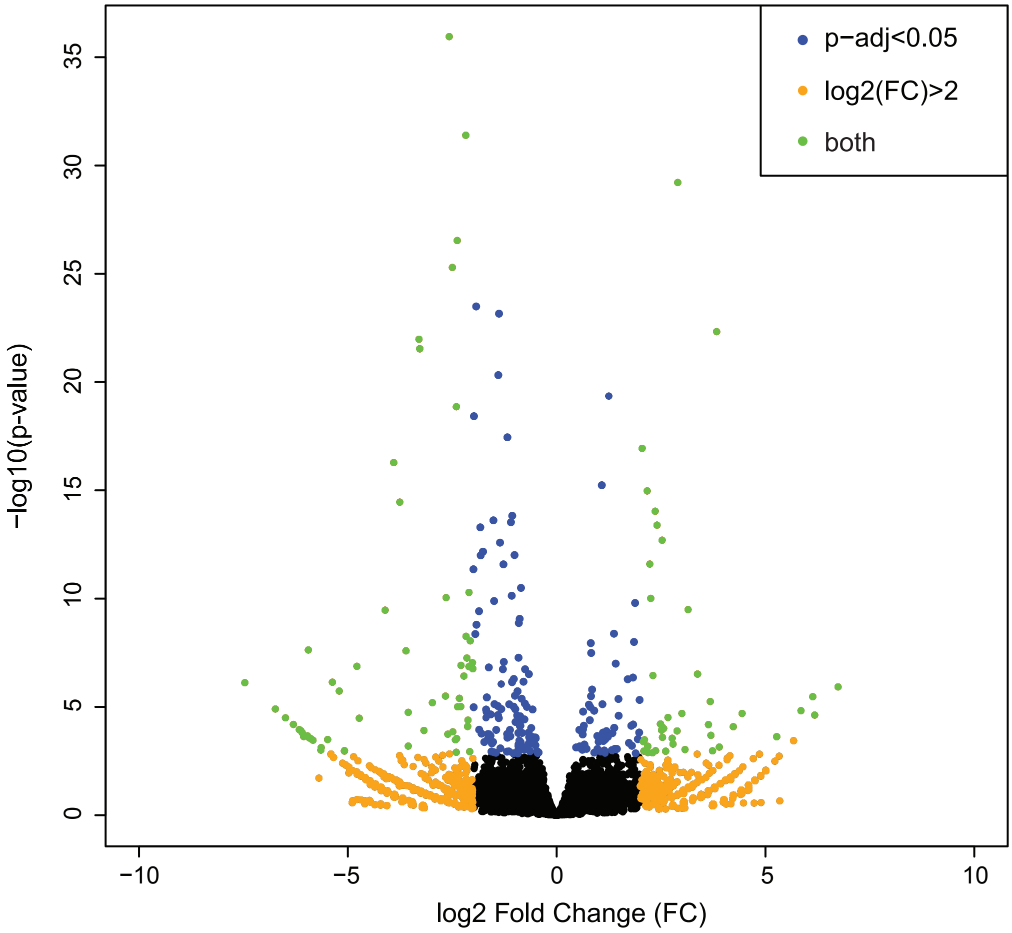


**Figure S4: Differential transcript isoform expression.** Volcano plot of differential isoform expression between undifferentiated and differentiated SH-SY5Y cells. An adjusted p-value of <0.05 from DESeq2 (green and blue dots) was considered significant for differential expression.
